# Parallel evolution of urban-rural clines in melanism in a widespread mammal

**DOI:** 10.1101/2021.09.08.459478

**Authors:** Bradley J. Cosentino, James P. Gibbs

## Abstract

Urbanization is the dominant trend of global land use change. The replicated nature of environmental change associated with urbanization should drive parallel evolution, yet insight into the repeatability of evolutionary processes in urban areas has been limited by a lack of multi-city studies. Here we leverage community science data on coat color in >60,000 eastern gray squirrels (*Sciurus carolinensis*) across 43 North American cities to test for parallel clines in melanism, a genetically based trait associated with thermoregulation and crypsis. We show the prevalence of melanism was positively associated with urbanization as measured by impervious cover. Urban-rural clines in melanism were strongest in the largest cities with extensive forest cover and weakest or absent in cities with warmer winter temperatures, where thermal selection likely limits the prevalence of melanism. Our results suggest that novel traits can evolve in a highly repeatable manner among urban areas, modified by factors intrinsic to individual cities, including their size, land cover, and climate.

## Introduction

Urban land use is expanding at a rate faster than urban population growth^1^, transforming land cover, climate, hydrology, and biogeochemical cycling^2^. The dramatic environmental change associated with urbanization renders cities similar to one another in many abiotic and biotic characteristics^3,4^. For example, cities have extensive impervious cover (e.g., buildings, transportation networks), increased air temperature, and increased air, water, light, and noise pollution relative to their adjacent rural landscapes^2^.

The convergence of environmental conditions in cities makes urban ecosystems an ideal focus for addressing longstanding questions in evolutionary biology about the degree to which populations exhibit parallel evolution, that is, when multiple populations occupying similar environments evolve comparable traits^5^. Cities are thought to drive parallel evolution as urban populations adapt to similar selection pressures^6,7,8,9^. Cities are not homogenous, however, and variation in their characteristics likely mediates processes that facilitate or inhibit parallel evolution, generating variation in the shape of genotypic or phenotypic clines along urbanization gradients. Clines may be strongest in large cities where the environmental gradients driving divergent selection between urban and rural areas are steepest. Conversely, clines may be weak or nonexistent in small cities or cities with extensive greenspace connecting urban and rural populations via gene flow. Environmental conditions at large spatial scales – such as regional climate – may also modulate the evolution of urban-rural clines at a smaller scale. For example, directional selection for traits that reduce thermal stress at climate extremes may overwhelm selective pressures associated with local land use, weakening urban-rural clines^10^. Advances in our understanding of how landscape-scale differences among cities facilitate or constrain parallel evolution have been limited by a lack of studies conducted across multiple, replicate cities^7,8,9^. Addressing this knowledge gap will help clarify how urbanization, the megatrend of global land use change, will shape biodiversity.

Here we test for urban-rural clines as a signal of parallel evolution and examine how landscape features mediate the shape of clines. Our focus is on melanism in eastern gray squirrels (*Sciurus carolinensis*) in 43 cities in North America (Fig. 1, Supplementary Fig. 1). Gray squirrels are among the most ubiquitous of fauna in cities and rural woodlands in eastern North America^11^. Coat color in gray squirrels is typically gray or black (melanic) and inherited in a simple Mendelian fashion at the melanocortin 1 receptor gene (*Mc1R*)^12^. Evidence suggests divergent selection associated with crypsis, predation risk, and road mortality favors the melanic morph in cities and the gray morph in rural woodlands^13^, which should cause a clinal decrease in melanism from urban to rural areas. However, urban-rural divergence in melanism has been documented in only two cities (Syracuse, NY^13^ and Wooster, OH^14^).

**Fig. 1.**
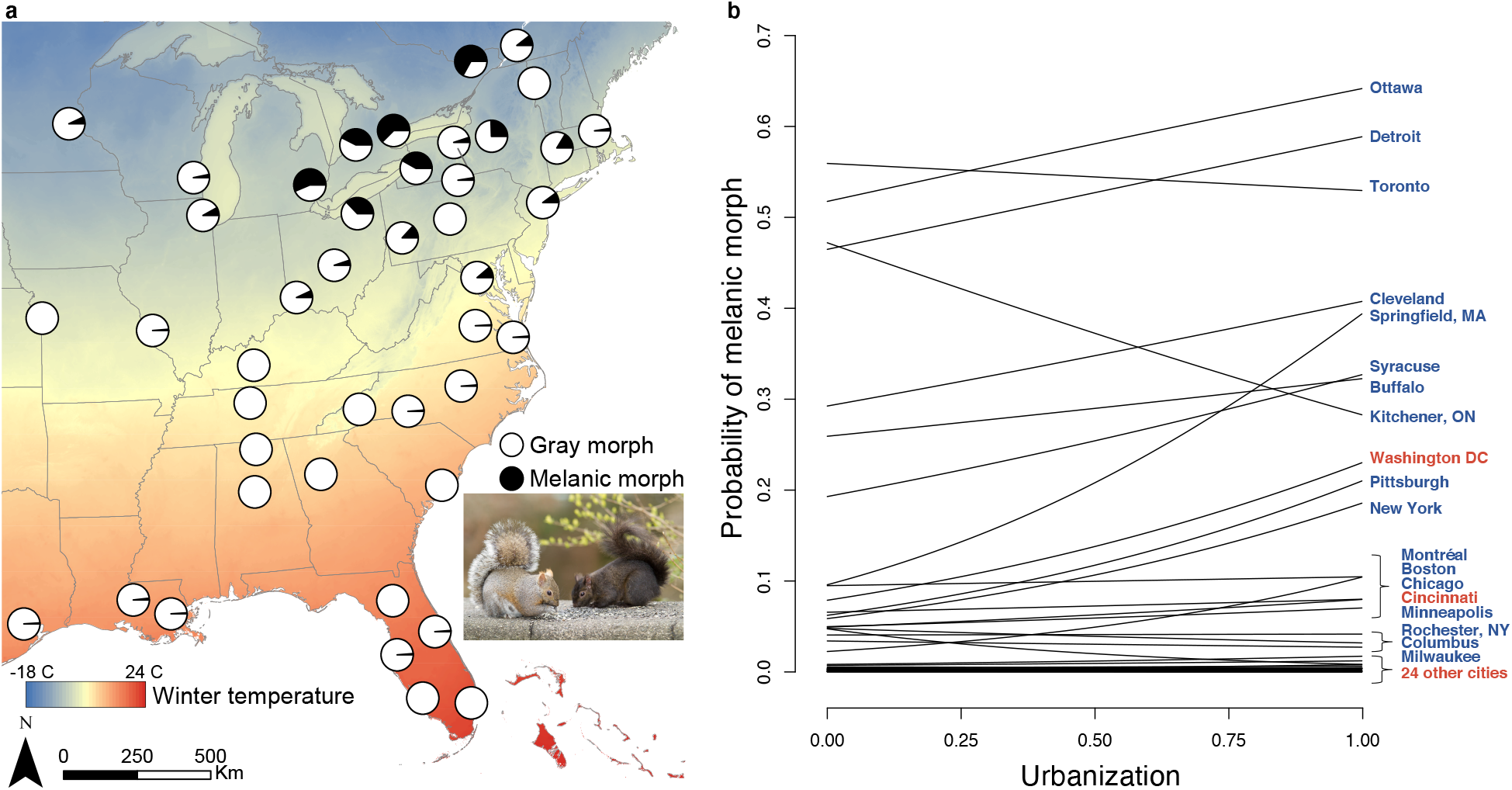
**a**, Distribution of coat color morphs of 26,924 eastern gray squirrels (*Sciurus carolinensis*) among 43 cities in North America. Pie charts show the proportion of melanic and gray color morphs in each city. Gray morph included rare color morphs that accounted for 2% of observations (Supplementary Table 1). **b**, Clines in melanism along urbanization gradients in each city. Urbanization was measured as standardized impervious cover. Regression lines represent predicted effect of urbanization on melanism in each city based on a linear mixed model including effects of city size, forest cover, and winter temperature (Fig. 2, Supplementary Table 2). City names are color-coded to show winter temperatures above (red) and below (blue) the median winter temperature. Photo ‘grey squirrel, black squirrel’ (tinyurl.com/pauha835) by Eyesplash Photography is licensed under CC BY-NC-ND 2.0 (https://creativecommons.org/licenses/by-nc-nd/2.0/).

We used a dataset on coat color from squirrel observations collected via the community science project *SquirrelMapper*^13^ to test for parallel clines in melanism along gradients of impervious cover. We included city size, forest cover across each urbanization gradient, and winter temperature in a linear mixed model to test how these landscape-scale characteristics affect the prevalence of melanism in each city (i.e., main effects of each covariate) and drive variation in cline shape among cities (i.e., interaction effects with impervious cover). Gray squirrels can disperse long distances (>10 km)^15^, but movements are limited by forest fragmentation^16^, so we expected clines to be weakest in small cities and cities with extensive forest cover where gene flow between urban and rural areas should be greatest. We included winter temperature in the model because melanic squirrels are known to have greater thermogenic capacity than the gray morph at cold temperature^17^. Clines may be weak in cities with cold winter temperature if thermal selection overwhelms the selection pressures on melanism associated with urbanization.

## Results and Discussion

Our analyses revealed that a positive relationship between the prevalence of melanism and degree of urbanization was replicated across multiple cities in North America (main effect of impervious cover = 0.17, SE = 0.04, *P* < 0.001; Figs. 1–2; Supplementary Table 2). Repeated shifts in phenotype along environmental gradients – such as the general increase in melanism from rural to urban areas documented here – is a hallmark of parallel evolution. Melanism in gray squirrels is a simple Mendelian trait coded by a 24-bp deletion at *Mc1R*^12^. Individuals with at least one copy of the deletion allele are melanic, so clines in melanism very likely occur at both phenotypic and genotypic levels.

**Fig. 2.**
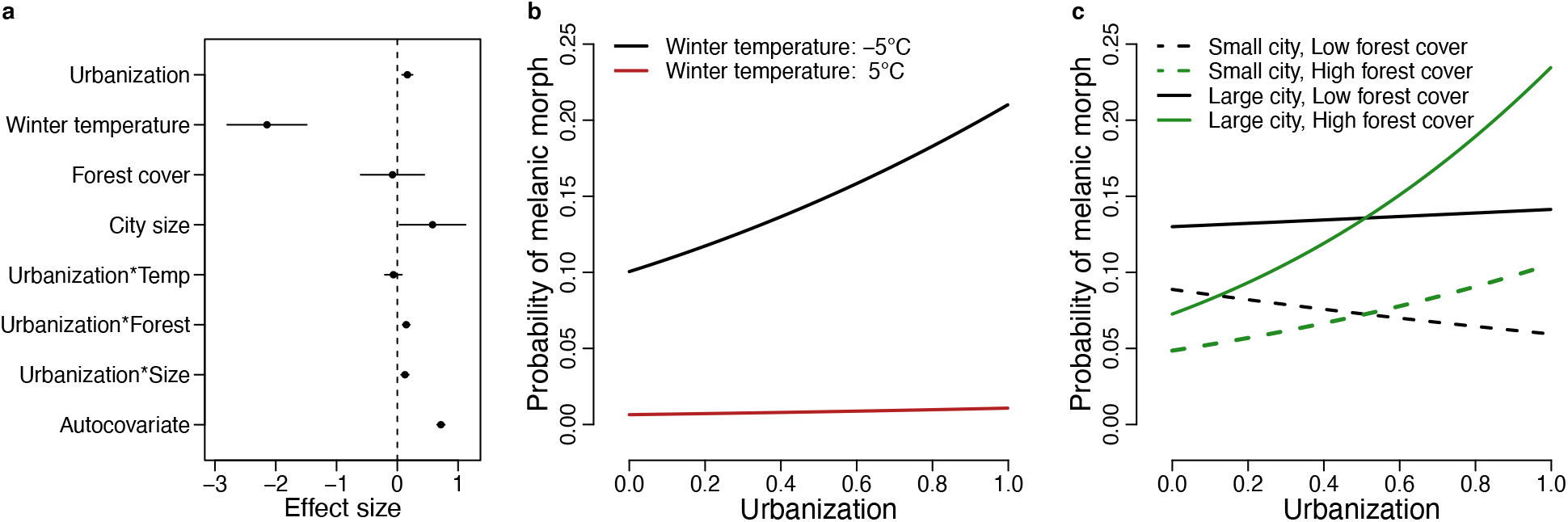
**a**, Effect sizes for fixed effects of urbanization, winter temperature, forest cover, city size, interaction terms with urbanization, and a spatial autocovariate from a linear mixed model of melanism in 26,924 eastern gray squirrels (*Sciurus carolinensis*) among 43 cities in North America. Effect sizes represent standardized regression coefficients and include 95% confidence intervals. **b**, Relationship of the probability of melanism to urbanization at low (−5°C) and high (5°C) winter temperature while holding city size and forest cover constant. **c**, Relationship of the probability of melanism to urbanization at varying levels of city size and forest cover. Effects of urbanization on melanism are shown for combinations of small cities (25,000 ha), large cities (350,000 ha), low forest cover (15%), and high forest cover (45%) while holding winter temperature constant at −5°C. Urbanization was measured as standardized impervious cover in all panels.

The presence of parallel clines in heritable traits is often considered evidence of adaptive evolution because gene flow should homogenize populations in the absence of natural selection^18^. Known drivers of parallel adaptive evolution in cities include the urban heat island^19^ and pollution^20^. Evidence suggests multiple selective mechanisms involving crypsis help maintain clines in melanism in gray squirrels. Predation – including hunting of squirrels by humans – is often the main driver of squirrel mortality in rural woodlands^21,22,23^, and the gray morph is more concealed than the melanic morph in regrown, secondary forests that dominate rural woodlands in our study area^13^. In contrast, predation pressure on tree squirrels tends to be relaxed in cities^21,22^, and relaxed selection against melanic squirrels in cities could generate the observed urban-rural clines. On the other hand, novel selection pressures in cities may favor melanics. The primary driver of mortality of tree squirrels in cities tends to be vehicular collisions^22^. The melanic morph is more visible to vehicle drivers than the gray morph on asphalt roads, which may lead to the melanic morph having greater survival in areas with high car traffic^13^. Additional selective mechanisms may play a role in maintaining urban melanism (e.g., parasite pressure, pollution)^24^, including mechanisms involving correlated traits^25^. For example, melanic squirrels have less phylogenetically variable microbiomes than the gray morph in cities^26^. A more complete picture of the selective mechanisms driving urban-rural clines will require linking trait variation to fitness differences between squirrel color morphs in urban and rural areas.

An alternative explanation for the urban-rural clines documented here is that melanic squirrels have been introduced by humans to some cities^23^. Strong genetic drift via founder events, combined with restricted gene flow along the urbanization gradient, can generate single-locus clines^27^. Variation in the size of founding populations likely causes some variation in the prevalence of melanism among cities, and thus the strength of urban-rural clines. At an extreme, urban-rural clines are absent in cities where melanism was historically absent and melanic squirrels have never been introduced. However, two factors lead us to believe that urban founder events are not the driving force behind the observed urban-rural clines. First, single-locus clines generated by neutral evolutionary forces require strong isolation-by-distance (IBD)^27^. We suspect IBD between urban and rural areas exists but is limited in strength by high effective population size in cities (e.g., urban squirrel densities 2.5 times greater than rural squirrel densities^28^), high capacity for long-distance dispersal^15^, and frequent translocation of squirrels^29^. Second, at northern latitudes where we observe most urban-rural clines, melanism was common in rural old growth forests prior to the 1800s^30,31,32^, including regions where it is now rare except in cities (e.g., upstate New York^13,31^). Founder events in cities do not explain the paucity of melanics in rural areas relative to the past. We hypothesize that the dramatic shift from old growth to secondary forests following European settlement in North America^33,34,35^ changed the degree to which gray and melanic morphs are concealed from visual predators and hunters, favoring the gray morph in regrown forests^13^. Genomic analysis and field experiments are needed to clarify the role of natural selection versus nonadaptive processes in generating the observed urban-rural clines^36^.

Although clines in melanism tended to take the same form among cities, the strength of urban-rural clines varied, supporting the idea that repeated evolution is often inconsistent^5^. First, there was a strong negative relationship between melanism and winter temperature (main effect of winter temperature = −2.14, SE = 0.33, *P* < 0.001; Figs. 1–2; Supplementary Table 2) that likely constrains the development of urban-rural clines. Urban-rural clines in melanism were most common in northern cities with cold winter temperature (Fig. 1b), whereas clines were absent in cities with warm winters where melanism was rare or nonexistent. The melanic morph has greater capacity for non-shivering thermogenesis and retains more heat than the gray morph at cold temperature^17,37,38^. Our results suggest thermal selection at a regional scale plays an important role in maintaining melanism in the north, allowing other processes operating at a finer spatial scale to cause the evolution of urban-rural clines. Winter temperature has also been shown to constrain the evolution of urban-rural clines in cyanogenesis in white clover (*Trifolium repens*)^39^. Importantly, our results do not suggest urban-rural clines in melanism in gray squirrels are caused by the urban heat island effect, which should cause the opposite clinal pattern than that observed.

Second, the positive relationship between urbanization and melanism was strongest in large cities with extensive forest cover (impervious*city size interaction effect = 0.13, SE = 0.03, *P* < 0.001; impervious*forest cover interaction effect = 0.15, SE = 0.03, *P* < 0.001; Fig. 2; Supplementary Table 2), where melanism increased by nearly 20% from rural to urban areas (Fig. 2c). In contrast, clines were weak to nonexistent in smaller cities and those with less extensive forest cover (Fig. 2c). Although gray squirrels have substantial capacity for long-distance dispersal, gene flow between urban and rural areas should be most limited in large cities due to isolation-by-distance and mortality caused by extensive road networks during dispersal.

Gene flow may be strongest between urban and rural populations in smaller cities, maintaining maladapted morphs across the urbanization gradient^9^. Selection associated with road mortality in urban areas may also be weak in small cities with limited traffic, which could explain the lower prevalence of melanism in smaller cities (main effect of city size = 0.58, SE = 0.28, *P* = 0.037; Fig. 2). Mechanistic studies are needed to provide insight into how road networks, traffic volume, and road-crossing structures (e.g., aerial utility lines that squirrels use to cross roads^23^) contribute to variation in gene flow and selection along urbanization gradients within and among cities.

The finding of strong clines in cities with extensive forest cover (Fig. 2) did not support our prediction that forest cover facilitates gene flow and weakens clines along the urbanization gradient. Gray squirrels require trees, and squirrel abundance and distribution along urbanization gradients is limited by tree cover^28^, particularly in large cities^40^. It is possible that cities with more extensive forest cover support a greater abundance of gray squirrels, increasing effective population size and thereby making selection more efficient in overcoming the effects of genetic drift on morph frequencies.

Community science datasets in ecology and evolution are powerful for conducting studies at large spatial scales and with levels of replication that would otherwise be infeasible. However, one challenge of using community science datasets is the potential for sampling biases^41^. For example, *SquirrelMapper* observers may be more likely to record observations of rare color morphs. Melanics were less common than the gray morph in most cities, so detection bias for rare morphs would inflate the estimated prevalence of melanism. However, we are unaware of a mechanism that would cause sampling bias to vary between urban and rural areas in a way that artificially generates the urban-rural clines documented here (e.g., bias toward recording the melanic morph in urban areas and gray morph in rural areas), so we have no reason to believe the relative change in melanism along urbanization gradients is biased.

Our work reinforces the idea that ecological homogenization among cities likely drives parallel evolution^19,20^, and it suggests cities can function as refuges for unique phenotypic traits^42^. Our study also provides insight into how landscape-scale differences among cities alter phenotypic clines along urbanization gradients. We show the degree of parallelism in urban-rural clines among cities is imperfect, with environmental heterogeneity among cities contributing to variation in the shape of clines. Greenspace availability in particular has been a focus in the ecological literature for understanding among-city variation in the response of species distributions to urbanization^40^ – our results show that greenspace availability in the form of forest cover likely mediates evolutionary responses to urbanization as well. Additional multi-city studies will be essential for understanding how the degree of parallel evolution depends on factors intrinsic to individual cities, including those not examined here (e.g., forest community composition). Finally, our work underscores the value of leveraging community science data to conduct studies of urban evolution across multiple cities.

## Methods

### Squirrel datasets

Squirrel records were sourced from the community science program *SquirrelMapper* (squirrelmapper.org). Starting in 2010, participants used a web platform to submit sightings of *S. carolinensis*, including the date of observation and color morph of each squirrel (“gray”, “melanic”, or “other”). Spatial coordinates for each observation were secured via Google Maps JavaScript API. A total of 5,899 records were submitted to the original

*SquirrelMapper* program from the native range of *S. carolinensis* in North America. Beginning in 2019, *SquirrelMapper* was integrated with *iNaturalist*, enabling users to submit georeferenced photos of each squirrel observation. All images of *S. carolinensis* submitted to *iNaturalist* through 21 December 2020 and classified as “research grade” were included for analysis (n = 75,130). Data from the original *SquirrelMapper* program were merged with the *iNaturalist* dataset for analyses.

We used the participatory science portal *Zooniverse* to crowdsource the classification of coat color of squirrels in images submitted to *iNaturalist*. Participants were asked to classify the number of squirrels observed in each image (“zero”, “one”, “two or more”) and the coat color of squirrels present in each image (“gray”, “melanic”, “other”, or “unclear”). Each image was classified by a minimum of 10 participants (range: 10-291; median = 10). We retained images with a minimum agreement threshold of 80% for the number of squirrels observed. Of the 72,239 images (96%) that met the agreement threshold, 865 were classified as having no squirrel present (e.g., track or other sign), 70,258 had a single squirrel, and 1,116 had two or more squirrels. Of the images classified as having at least one squirrel, we retained 63,506 images that had a minimum agreement threshold of 80% for coat color of at least one squirrel in the image. We removed 5,182 images that did not have location data available and 4,035 images taken outside of North America where *S. carolinensis* is introduced. The final *iNaturalist* dataset consisted of 54,289 images, of which 54,215 were classified as having a single coat color present (gray: 46,630, melanic: 6,728, other: 857) and 74 had two color morphs present (gray and melanic: 68, gray and other: 5, melanic and other: 1). A random sample of 500 images classified by the authors showed that the classification scheme we used with 80% agreement thresholds was 100% accurate for the number of squirrels observed in the image and 99.8% accurate for coat color, the only discrepancy being a single image classified as melanic that the authors classified as unclear. The final *iNaturalist* dataset was merged with squirrel records from the original *SquirrelMapper* project, yielding 60,262 squirrel records in total. Multiple coat colors observed at the same location were treated as separate records in this final dataset. Observations in the final dataset were made between 1980 and 2020 (median = 2019), with 97% of observations being reported since 2010. The 3% of observations prior to 2010 consisted of *iNaturalist* records and records by the authors and colleagues prior to *SquirrelMapper*.

### City selection

We selected cities in the native range of *S. carolinensis* that had at least 100 squirrel observations in the city and surrounding rural area. We defined the spatial extent of each city’s footprint using the Urban Area National Shapefile Record Layout^43^ for the United States and the Population Center Boundary File^44^ for Canada. We calculated the distance from the centroid of each city footprint to its maximum extent in each of the four cardinal directions. We then buffered each urban area by 25% of the average distance from the city centroid to its extent (Supplementary Fig. 2). This buffer distance was large enough to include a substantial area of rural land surrounding each city but not so large as to produce excessive overlap between polygons of adjacent cities. We determined the number of squirrels in each city polygon (encompassing the city footprint and hinterland) after spatially thinning the squirrel records to a minimum distance of 10 m using the *spThin* package (v. 0.2.0)^45^ in R^46^, which helps minimize spatial sampling bias. If two cities with ≥100 squirrels had polygons that overlapped, we retained the city with a greater sample size or spatial spread of squirrel observations. The final dataset included 26,924 squirrels among 43 cities (Fig. 1), and the median number of squirrels in each city was 254 (range: 101 – 4,731; Supplementary Table 1; maps of squirrel observations in each city available for download at https://github.com/bcosentino/squirrelMapper-urban-rural-clines).

### Environmental covariates

We measured percent impervious cover within 1 km around each squirrel observation using the Global Man-made Impervious Surface Dataset (30-m resolution)^47^. A 1-km buffer was chosen to encompass the typical home range size of *S. carolinensis* (<5 ha) and long-distance movements typical of males during winter and summer mating seasons^48^. For each city and its surrounding rural area we measured forest cover and winter temperature. Forest cover was measured using the Global 2010 Tree Cover database (30-m resolution)^49^, and we used WorldClim (version 2.1) to measure winter temperature as mean temperature of the coldest quarter (BIO11, 30s / 1 km resolution)^50^. We also measured the area of each city footprint as a metric of city size. All spatial analyses were conducted in R using the *raster* (v. 3.3-13)^51^ and *exactextractr* packages (v. 0.5.1)^52^.

### Statistical analyses

We fit a generalized linear mixed model with a binomial error distribution to test for effects of impervious cover, city size, winter temperature, and forest cover on the distribution of melanism. Melanism was the response variable and represented by a binary indicator, with 1 corresponding to melanic and 0 to other coat colors for each squirrel observation. Impervious cover within 1 km of each observation was included as a fixed effect to represent each city’s urbanization gradient. Because the range of impervious cover varied among cities, we rescaled impervious cover within each city to range from 0 (lowest impervious cover) to 1 (greatest impervious cover)^39^. This ensured the slope term for impervious cover has a consistent interpretation among cities, allowing us to test for a general urban-to-rural cline in melanism. City size (log-transformed), forest cover, and winter temperature were included as fixed effects to test whether these variables predict variation in the prevalence of melanism among cities. Two-way interaction terms were included between each predictor and impervious cover to test how city characteristics affect the shape of urban-to-rural clines in melanism. We standardized each predictor so that each variable had mean = 0 and SD = 1, which aided in model convergence and allowed us to compare the effect size of each slope term^53^. We also included city as a random effect to account for the nested observations within each city and to estimate unexplained variance in the prevalence of melanism among cities (Supplementary Table 3). The model was fit with the *lme4* package in R (v. 1.1-26)^54^. Two-sided Wald tests were conducted for each parameter in the model. Binned residual plots were generated using the *arm* package (v. 1.11-2)^55^ as recommended for models with a binary response variable^56^. Plots of binned residuals on fitted values and each predictor suggested adequate model fit and no substantial deviation from model assumptions.

We used the residual autocovariate approach^57^ to account for spatial autocorrelation in the residuals of the model. We extracted residuals from the global model and created a spatial autocovariate for the residuals using the *spdep* package (v. 1.1-7)^58^. We set the radius neighborhood to 31 km (the greatest nearest neighbor distance in the thinned dataset) and used an inverse distance weighting scheme with style “B” for symmetric weights^59^. Our global model was then refit with the spatial autocovariate term as an additional predictor, which reduced spatial autocorrelation in the residuals from a maximum of *r* = 0.11 within 1 km to *r* < 0.05 (Moran’s *I*; Supplementary Fig. 3).

We refit our model using different spatial thinning distances (10 m, 50 m, 100 m) and buffer distances to measure impervious cover (500 m, 1 km, 10 km). These analyses showed the model results were not sensitive to spatial thinning distances (Supplementary Table 4) or buffer distances to quantify impervious cover around squirrel observations (Supplementary Table 5).

## Acknowledgements

This research was supported by the National Science Foundation (DEB 2018249). We thank B. Fischman for comments on the manuscript.

## Author Contributions

B.J.C. and J.P.G designed the study. B.J.C. led the data analysis and wrote the first draft of the manuscript. J.P.G. contributed ideas for data analysis and edited the manuscript.

## Data Availability

The observations of eastern gray squirrels (*Sciurus carolinensis*) generated and analyzed for this project can be found at https://github.com/bcosentino/squirrelMapper-urban-rural-clines.

