## Supplemental Information for "Parallel evolution of urban-rural clines in melanism in a widespread mammal"

This file includes:

Supplementary Table 1: Number of *Sciurus carolinensis* observations in total and by color morph in each city selected for analysis.

Supplementary Table 2: Parameter estimates for fixed effects in a linear mixed model of the probability of melanism in eastern gray squirrels (*Sciurus carolinensis*).

Supplementary Table 3: Parameter estimates for the random effect of city in a linear mixed model of the probability of melanism in eastern gray squirrels (*Sciurus carolinensis*). The estimated standard deviation for the random effect among cities was 1.68.

Supplementary Table 4: Parameter estimates for fixed effects in a linear mixed model of the probability of melanism in eastern gray squirrels (*Sciurus carolinensis*) using alternative spatial thinning distances (10 m, 50 m, 100 m).

Supplementary Table 5: Parameter estimates for fixed effects in a linear mixed model of the probability of melanism in eastern gray squirrels (*Sciurus carolinensis*) using alternative buffer distances to quantify impervious cover around each squirrel observation (500 m, 1 km, 10 km).

Supplementary Figure 1. Map of 43 cities in North America included in analyses of the distribution of coat color morphs of eastern gray squirrels (*Sciurus carolinensis*).

Supplementary Figure 2. Map of Syracuse, NY highlighting methodology for defining the spatial extent of the city footprint and buffer.

Supplementary Figure 3. Spatial correlograms of residuals from the linear mixed model of the probability of melanism. Moran's *I* is shown for distance classes of 1 km for a model either without or with a spatial autocovariate included in the model.

**Supplementary Table 1.** Number of *Sciurus carolinensis* observations in total and by color morph in each city selected for analysis.

| City name | *S. carolinensis* observations | Gray morph | Melanic morph | Other morphs |
|---|---|---|---|---|
| Asheville, NC | 164 | 142 | 0 | 22 |
| Atlanta, GA | 509 | 506 | 0 | 3 |
| Baton Rouge, LA | 177 | 171 | 3 | 3 |
| Birmingham, AL | 106 | 105 | 0 | 1 |
| Boston, MA | 3104 | 2971 | 84 | 49 |
| Bowling Green, KY | 164 | 108 | 0 | 56 |
| Buffalo, NY | 203 | 114 | 85 | 4 |
| Burlington, VT | 197 | 196 | 0 | 1 |
| Cape Coral, FL | 107 | 107 | 0 | 0 |
| Charleston, SC | 129 | 129 | 0 | 0 |
| Charlotte, NC | 237 | 233 | 3 | 1 |
| Chicago, IL | 1047 | 962 | 84 | 1 |
| Cincinnati, OH | 732 | 668 | 50 | 14 |
| Cleveland, OH | 298 | 183 | 111 | 4 |
| Columbus, OH | 332 | 281 | 17 | 34 |
| Detroit, MI | 282 | 116 | 159 | 7 |
| Gainesville, FL | 254 | 254 | 0 | 0 |
| Geneva, NY | 117 | 113 | 3 | 1 |
| Houston, TX | 885 | 878 | 4 | 3 |
| Huntsville, AL | 178 | 177 | 0 | 1 |
| Kansas City, MO | 101 | 98 | 0 | 3 |
| Kitchener, Ontario | 200 | 114 | 85 | 1 |
| Miami, FL | 777 | 774 | 0 | 3 |
| Milwaukee, WI | 346 | 328 | 11 | 7 |
| Minneapolis, MN | 570 | 426 | 40 | 104 |
| Montréal, Quebec | 770 | 648 | 80 | 42 |
| Nashville, TN | 117 | 116 | 0 | 1 |
| New Orleans, LA | 118 | 117 | 1 | 0 |
| New York, NY | 4731 | 4236 | 480 | 15 |
| Orlando, FL | 459 | 451 | 1 | 7 |
| Ottawa, Ontario | 697 | 230 | 466 | 1 |
| Pittsburgh, PA | 226 | 193 | 30 | 3 |
| Raleigh, NC | 681 | 668 | 1 | 12 |
| Richmond, VA | 267 | 258 | 1 | 8 |
| Rochester, NY | 122 | 113 | 6 | 3 |
| Springfield, MA | 140 | 117 | 23 | 0 |
| St. Louis, MO | 253 | 250 | 1 | 2 |
| State College, PA | 253 | 253 | 0 | 0 |
| Syracuse, NY | 736 | 546 | 189 | 1 |
| Tampa, FL | 1520 | 1518 | 1 | 1 |
| Toronto, Ontario | 1723 | 615 | 1075 | 33 |
| Virginia Beach, VA | 251 | 249 | 1 | 1 |
| Washington DC | 2644 | 2271 | 298 | 75 |

**Supplementary Table 2.** Parameter estimates for fixed effects in a linear mixed model of the probability of melanism in eastern gray squirrels (*Sciurus carolinensis*).

| Fixed effect | Estimate | SE | *z* | *P* |
|---|---|---|---|---|
| Intercept | -3.73 | 0.35 | -10.67 | <0.001 |
| Impervious cover | 0.17 | 0.04 | 3.76 | <0.001 |
| Winter temperature | -2.14 | 0.33 | -6.43 | <0.001 |
| Forest cover | -0.08 | 0.27 | -0.30 | 0.766 |
| City size | 0.58 | 0.28 | 2.09 | 0.037 |
| Impervious*Temperature | -0.07 | 0.07 | -0.91 | 0.363 |
| Impervious*Forest cover | 0.15 | 0.03 | 4.92 | <0.001 |
| Impervious*City size | 0.13 | 0.03 | 3.91 | <0.001 |
| Spatial autocovariate | 0.72 | 0.03 | 22.28 | <0.001 |

**Supplementary Table 3.** Parameter estimates for the random effect of city in a linear mixed model of the probability of melanism in eastern gray squirrels (*Sciurus carolinensis*). The estimated standard deviation for the random effect among cities was 1.68.

| City name | Conditional modes |
|---|---|
| Asheville, NC | -1.20 |
| Atlanta, GA | -2.17 |
| Baton Rouge, LA | 3.19 |
| Birmingham, AL | -0.57 |
| Boston, MA | -0.98 |
| Bowling Green, KY | -0.62 |
| Buffalo, NY | 2.34 |
| Burlington, VT | -2.98 |
| Cape Coral, FL | -0.05 |
| Charleston, SC | -0.38 |
| Charlotte, NC | 0.85 |
| Chicago, IL | -1.14 |
| Cincinnati, OH | 0.79 |
| Cleveland, OH | 2.60 |
| Columbus, OH | 0.09 |
| Detroit, MI | 2.57 |
| Gainesville, FL | -0.14 |
| Geneva, NY | 1.25 |
| Houston, TX | 1.05 |
| Huntsville, AL | -1.00 |
| Kansas City, MO | -2.33 |
| Kitchener, Ontario | 2.91 |
| Miami, FL | -0.33 |
| Milwaukee, WI | -1.72 |
| Minneapolis, MN | -2.68 |
| Montréal, Quebec | -1.38 |
| Nashville, TN | -1.34 |
| New Orleans, LA | 2.16 |
| New York, NY | 0.24 |
| Orlando, FL | 1.58 |
| Ottawa, Ontario | 2.06 |
| Pittsburgh, PA | 1.14 |
| Raleigh, NC | -1.16 |
| Richmond, VA | -0.55 |
| Rochester, NY | -0.23 |
| Springfield, MA | 1.94 |
| St. Louis, MO | -2.19 |
| State College, PA | -1.62 |
| Syracuse, NY | 2.37 |
| Tampa, FL | 0.59 |
| Toronto, Ontario | 2.77 |
| Virginia Beach, VA | -0.37 |
| Washington DC | 2.22 |

**Supplementary Table 4.** Parameter estimates for fixed effects in a linear mixed model of the probability of melanism in eastern gray squirrels (*Sciurus carolinensis*) using alternative spatial thinning distances (10 m, 50 m, 100 m). Cities included in each dataset had a minimum of 100 squirrel observations. Samples sizes are included for the number of cities and squirrel observations included in each dataset.

| Thinning distance | Cities | Squirrels | Fixed effect | Estimate | SE | *z* | *P* |
|---|---|---|---|---|---|---|---|
| 10 m | 43 | 26,924 | Intercept | -3.73 | 0.35 | -10.67 | <0.001 |
| | | | Impervious cover | 0.17 | 0.04 | 3.76 | <0.001 |
| | | | Winter temperature | -2.14 | 0.33 | -6.43 | <0.001 |
| | | | Forest cover | -0.08 | 0.27 | -0.30 | 0.766 |
| | | | City size | 0.58 | 0.28 | 2.09 | 0.037 |
| | | | Impervious*Temperature | -0.07 | 0.07 | -0.91 | 0.363 |
| | | | Impervious*Forest cover | 0.15 | 0.03 | 4.92 | <0.001 |
| | | | Impervious*City size | 0.13 | 0.03 | 3.91 | <0.001 |
| | | | Spatial autocovariate | 0.72 | 0.03 | 22.28 | <0.001 |
| 50 m | 38 | 20,285 | Intercept | -3.54 | 0.35 | -10.13 | <0.001 |
| | | | Impervious cover | 0.17 | 0.05 | 3.47 | <0.001 |
| | | | Winter temperature | -2.27 | 0.35 | -6.45 | <0.001 |
| | | | Forest cover | -0.15 | 0.29 | -0.52 | 0.600 |
| | | | City size | 0.60 | 0.32 | 1.91 | 0.056 |
| | | | Impervious*Temperature | -0.04 | 0.08 | -0.45 | 0.655 |
| | | | Impervious*Forest cover | 0.20 | 0.03 | 6.29 | <0.001 |
| | | | Impervious*City size | 0.13 | 0.03 | 3.91 | <0.001 |
| | | | Spatial autocovariate | 0.57 | 0.03 | 18.98 | <0.001 |
| 100 m | 34 | 16,991 | Intercept | -3.43 | 0.34 | -10.02 | <0.001 |
| | | | Impervious cover | 0.18 | 0.05 | 3.50 | <0.001 |
| | | | Winter temperature | -2.07 | 0.33 | -6.20 | <0.001 |
| | | | Forest cover | -0.07 | 0.27 | -0.26 | 0.797 |
| | | | City size | 0.30 | 0.30 | 1.00 | 0.318 |
| | | | Impervious*Temperature | -0.05 | 0.08 | -0.54 | 0.592 |
| | | | Impervious*Forest cover | 0.22 | 0.03 | 6.46 | <0.001 |
| | | | Impervious*City size | 0.13 | 0.03 | 3.86 | <0.001 |
| | | | Spatial autocovariate | 0.49 | 0.03 | 16.64 | <0.001 |

**Supplementary Table 5.** Parameter estimates for fixed effects in a linear mixed model of the probability of melanism in eastern gray squirrels (*Sciurus carolinensis*) using alternative buffer distances to quantify impervious cover around each squirrel observation (500 m, 1 km, 10 km).

| Impervious cover buffer distance | Fixed effect | Estimate | SE | *z* | *P* |
|---|---|---|---|---|---|
| 500 m | Intercept | -3.74 | 0.35 | -10.71 | <0.001 |
| | Impervious cover | 0.15 | 0.04 | 3.48 | <0.001 |
| | Winter temperature | -2.15 | 0.33 | -6.45 | <0.001 |
| | Forest cover | -0.09 | 0.27 | -0.32 | 0.748 |
| | City size | 0.60 | 0.28 | 2.17 | 0.030 |
| | Impervious*Temperature | 0.02 | 0.07 | 0.23 | 0.820 |
| | Impervious*Forest cover | 0.12 | 0.03 | 4.14 | <0.001 |
| | Impervious*City size | 0.08 | 0.03 | 2.36 | 0.018 |
| | Spatial autocovariate | 0.70 | 0.03 | 21.97 | <0.001 |
| 1 km | Intercept | -3.73 | 0.35 | -10.67 | <0.001 |
| | Impervious cover | 0.17 | 0.04 | 3.76 | <0.001 |
| | Winter temperature | -2.14 | 0.33 | -6.43 | <0.001 |
| | Forest cover | -0.08 | 0.27 | -0.30 | 0.766 |
| | City size | 0.58 | 0.28 | 2.09 | 0.037 |
| | Impervious*Temperature | -0.07 | 0.07 | -0.91 | 0.363 |
| | Impervious*Forest cover | 0.15 | 0.03 | 4.92 | <0.001 |
| | Impervious*City size | 0.13 | 0.03 | 3.91 | <0.001 |
| | Spatial autocovariate | 0.72 | 0.03 | 22.28 | <0.001 |
| 10 km | Intercept | -3.80 | 0.36 | -10.68 | <0.001 |
| | Impervious cover | 0.26 | 0.05 | 5.41 | <0.001 |
| | Winter temperature | -2.15 | 0.34 | -6.35 | <0.001 |
| | Forest cover | -0.10 | 0.27 | -0.38 | 0.708 |
| | City size | 0.58 | 0.28 | 2.06 | 0.040 |
| | Impervious*Temperature | -0.06 | 0.08 | -0.81 | 0.421 |
| | Impervious*Forest cover | 0.23 | 0.03 | 7.29 | <0.001 |
| | Impervious*City size | 0.23 | 0.04 | 6.27 | <0.001 |
| | Spatial autocovariate | 0.74 | 0.03 | 23.25 | <0.001 |

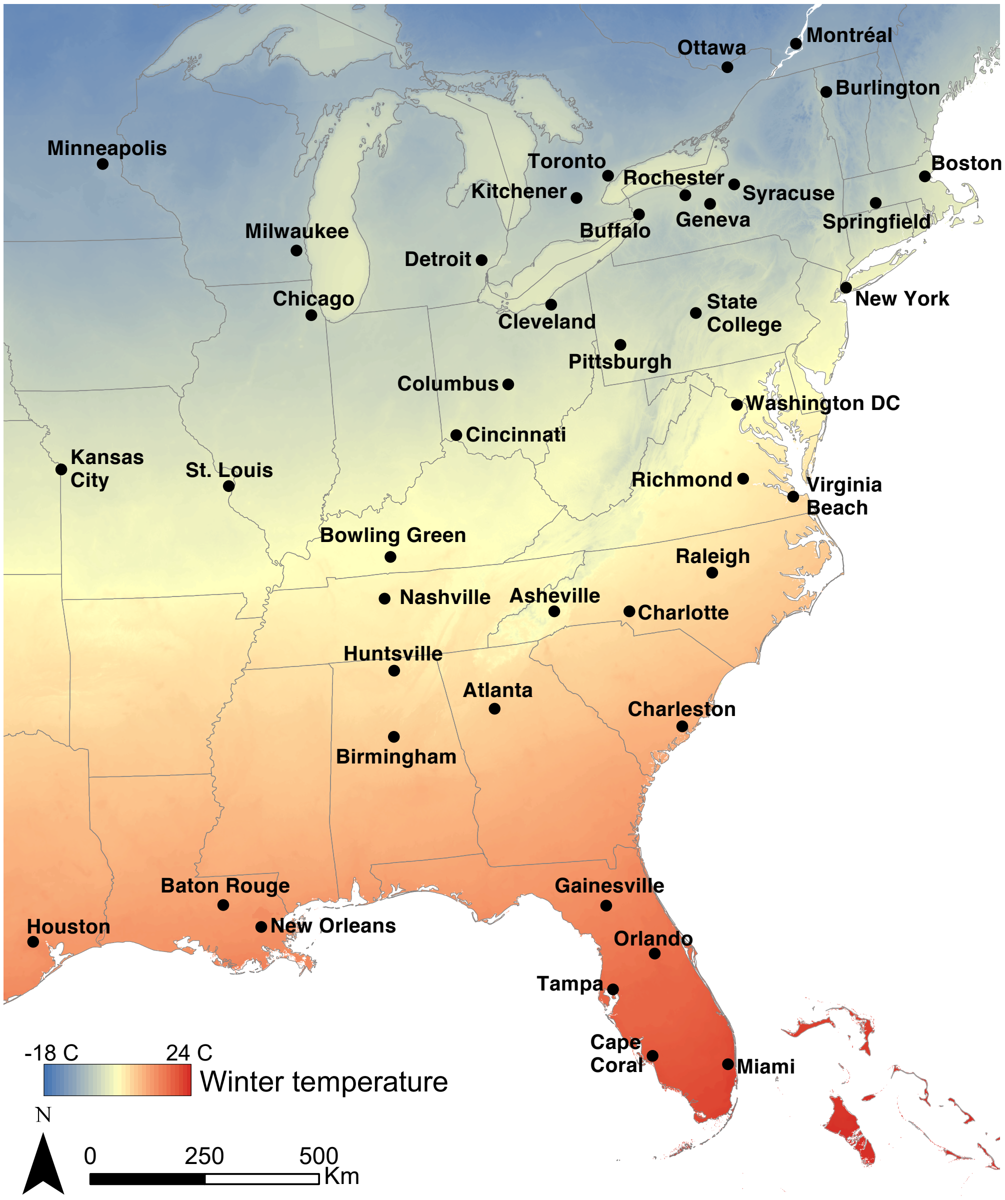


**Supplementary Figure 1.** Map of 43 cities in North America included in analyses of the distribution of coat color morphs of eastern gray squirrels (*Sciurus carolinensis*).

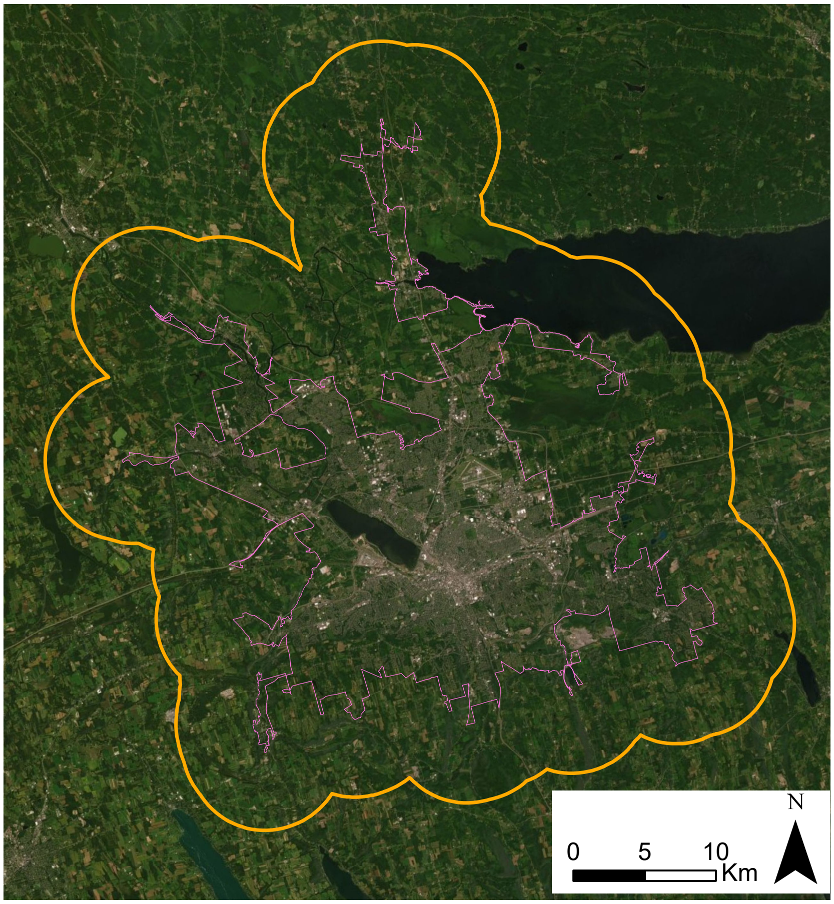


**Supplementary Figure 2.** Map of Syracuse, NY highlighting methodology for defining the spatial extent of the city footprint (purple line) and buffer (orange line).

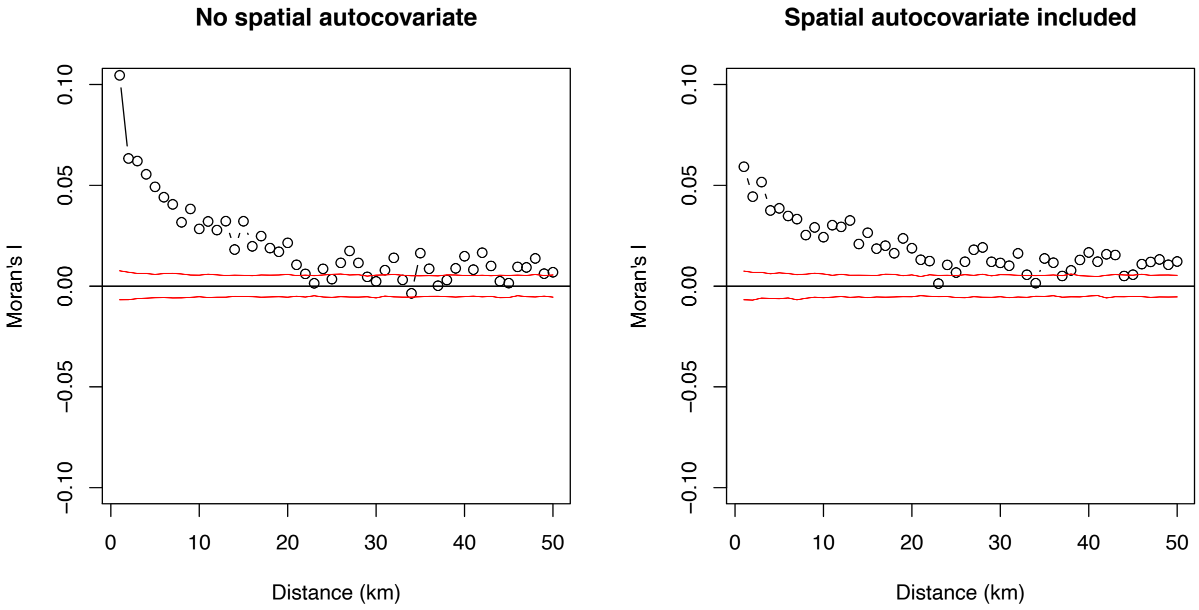


**Supplementary Figure 3.** Spatial correlograms of residuals from the linear mixed model of the probability of melanism (Supplementary Table 2). Moran's *I* is shown for distance classes of 1 km for a model either without or with a spatial autocovariate included in the model. Red lines show bounds of Moran's *I* beyond which the null hypothesis of $r = 0$ is rejected at a significance value of 0.05, based on Monte Carlo permutations with 999 permutations at each distance class. The bounds of statistical significance are conservative because the analysis does not account for the nesting of observations within cities.
